# Specific induction of double negative B cells during protective and pathogenic immune responses

**DOI:** 10.1101/2020.09.08.285148

**Authors:** Christoph Ruschil, Gisela Gabernet, Gildas Lepennetier, Simon Heumos, Miriam Kaminski, Zsuzsanna Hrasko, Martin Irmler, Johannes Beckers, Ulf Ziemann, Sven Nahnsen, Greg P. Owens, Jeffrey L. Bennett, Bernhard Hemmer, Markus C. Kowarik

**Affiliations:** Department of Neurology & Stroke, Eberhard-Karls University, Tübingen, Germany; Hertie Institute for Clinical Brain Research, Eberhard-Karls University, Tübingen, Germany; Quantitative Biology Center (QBiC), Eberhard-Karls University of Tübingen, Tübingen, Germany; Department of Neurology, Technische Universität München, Munich, Germany; Department of Psychiatry and Psychotherapy, Charite Universitätsmedizin Berlin, Germany; Department of Internal Medicine 1, Universitätsklinikum Erlangen, Friedrich-Alexander Universität Erlangen-Nürnberg, Erlangen, Germany; Institute of Experimental Genetics, Helmholtz Zentrum München GmbH, 85764 Neuherberg, Germany; German Center for Diabetes Research (DZD), 85764 Neuherberg, Germany; Technische Universität München, Chair of Experimental Genetics, 85354 Freising, Germany; Department of Biochemistry and Molecular Genetics, University of Colorado, 13001 East 17h Place, Aurora, CO 80045; Departments of Neurology and Ophthalmology, Programs in Neuroscience and Immunology University of Colorado School of Medicine, 12700 East 19th Avenue, Aurora, CO 80045; Munich Cluster for Systems Neurology (SyNergy), Feodor-Lynen-Str. 17, 81377 Munich, Germany

**Author notes:** **Correspondence:** Markus C. Kowarik.

**Keywords:** B cells, double negative B cells, vaccination, autoimmune disorders, influenza vaccination, TBE vaccination, NMOSD

## Abstract

Double negative (DN) (CD19^+^CD20^low^CD27^−^IgD^−^) B cells are expanded in patients with autoimmune and infectious diseases; however their role in the humoral immune response remains unclear. Using systematic flow cytometric analyses of peripheral blood B cell subsets, we observed an inflated DN B cell population in patients with variety of active inflammatory conditions: myasthenia gravis, Guillain-Barré syndrome, neuromyelitis optica spectrum disorder, meningitis/encephalitis, and rheumatic disorders. Furthermore, we were able to induce DN B cells in healthy subjects following vaccination against influenza and tick borne encephalitis virus. Transcriptome analysis revealed a gene expression profile in DN B cells that clustered with naïve B cells, memory B cells, and plasmablasts. Immunoglobulin VH transcriptome sequencing and analysis of recombinant antibodies revealed clonal expansion of DN B cells, that were targeted against the vaccine antigen. Our study suggests that DN B cells are expanded in multiple inflammatory neurologic diseases and represent an inducible B cell population that responds to antigenic stimulation, possibly through an extra-follicular maturation pathway.

## 2 Introduction

Circulating B cell subsets are grossly classified as naïve, memory B cells and antibody secreting B cells (plasmablasts and plasma cells) that are generated following established maturation pathways (1). Recently, novel B cell subsets, activated naïve B cells (2) and double negative (DN) B cells (3) have gathered attention due to their expansion and activation in autoimmune disorders such as systemic lupus erythematosus.

Elevated numbers of DN B cells are found in aged individuals (4,5) and a heterogeneous group of inflammatory and infectious conditions. We recently showed that CD27^−^IgD^−^CD20^low^ double negative (DN) B cells are elevated in the peripheral blood (PB) of patients with active neuromyelitis optica spectrum disorder (NMOSD) and are closely related to disease relevant aquaporin-4 specific CSF B cell clones (6). In multiple sclerosis (MS), DN B cells have been shown to be upregulated in around 30% of MS patients, and their expansion may be linked to immune aging (7) or specific disease stages (4,8,9). Besides systemic lupus erythematosus (SLE) (2,10–12), peripheral blood DN B cells have been shown to be elevated in rheumatoid arthritis (RA)(13,14), Hashimoto’s thyreoiditis (15), and inflammatory bowel disease (16). DN B cells are also expanded following viral infection (17–19), bacterial sepsis (20), active malaria (21,22), and immunogenic tumors such as non-small cell lung cancer (23).

In general, CD27 expression on B cells has been considered to be a hallmark for somatically (hyper-) mutated antigen experienced B cells (24–26). Although DN B cells do not express CD27, this population bears signatures of antigen experienced B cells in terms of surface phenotype, proliferation response and patterns of somatic hypermutations (11,24,27,28). Immunoglobulin heavy chain variable region (IGHV) gene usage of DN B cells is closely related to class-switched CD27^+^ memory B cells (29). In addition, a large portion of DN B cells show somatic hypermutations although the mutational load is slightly lower than in class switched memory B cells (7). The distinct expression profile of the anti-apoptotic molecule Bcl2 and absence of ATP-binding cassette B1 transporter (ABCB1) has been used to discriminate DN B cells from naïve CD27^−^ B cells (5,27). However, DN B cells appear to be a heterogenic B cell subset (30) and have been suggested to represent either exhausted memory B cells / senescent B cells (31), transient effector B cells or a unique atypical memory-like B cells, which may be relevant for plasmablast generation (32,33). More recently, detailed studies in SLE indicated that a subset of DN B cells might be derived from an activated naïve B cell subset and further differentiate into plasmablasts through an extra-follicular maturation pathway (1).

The purpose of this study was two-fold: first, to enumerate the spectrum of inflammatory and infectious disorders with expanded DN B cell populations and second, to further characterize the relationship of DN B cells to other B cell subsets under defined antigenic stimulation.

## 3 Materials and methods

### 3.1 Standard protocol approvals, registrations and patients

You may insert up to 5 heading levels into your manuscript as can be seen in “Styles” tab of this template. These formatting styles are meant as a guide, as long as the heading levels are clear, Frontiers style will be applied during typesetting. All patients and subjects were recruited at the Department of Neurology, Technische Universität München and consented to the scientific use of their biologic samples. The study was approved by the local ethics committee of the Technische Universität München. Patients with the diagnoses of inflammatory diseases including Guillain–Barré syndrome (GBS, n = 23; all active disease stage), rheumatic diseases (n = 12; SLE n = 3, granulomatosis with polyangiitis n = 3, Sjögren syndrome n = 2, scleroderma, dermatomyositis, antiphospholipid syndrome each n = 1), meningitis (n = 20; viral including herpes simplex virus, varicella-zoster virus, enterovirus, tick borne encephalitis virus n = 7 unknown virus n = 4, bacterial including listeria, tuberculosis, spirochete, pneumococcus n = 9), neuromyelitis optica spectrum disorder (NMOSD, n = 10; active disease stage n = 5), Myasthenia gravis (n = 11; active disease stage n = 7), multiple sclerosis (MS, n = 21; relapse n = 14; no relapse n = 7) and controls with non-inflammatory neurological diseases (NIND, n = 29; diagnoses: headache n = 13, paresthesia of unknown origin / somatoform disorders n = 9, idiopathic intracranial hypertension n = 2, others n = 5) were recruited between 2014 and 2016. Further patients characteristics are shown in Table 1.

**Table 1:** Clinical characteristics of patients with neuro-inflammatory diseases and controls.

| **Diseases** | **Gender**<br>**F / M** | **Age mean**<br>**(range)** | **Disease**<br>**duration**<br>**(range)** | **Clinical**<br>**scores**<br>**(range)** | **Therapy at time-point of**<br>**analysis** |
| --- | --- | --- | --- | --- | --- |
| **Non-inflammatory**<br>**neurological**<br>**controls (NIND)**<br>**n = 29** | 23 / 6 | 39.7 y<br>(18 – 76) | 20.4 d<br>(3 – 70) | N. A. | None |
| **Meningitis /**<br>**Encephalitis**<br>**n = 20** | 5 / 15 | 40.6 y<br>(19 – 71) | 30.5 d<br>(1 – 183) | N. A. | Antibiotics (*e.g. ampicillin,*<br>*ceftriaxon, doxycyclin,*<br>*meropenem, vancomycin,*<br>*cefalexin*) (n = 16)<br>Virostatica (Aciclovir) (n = 8) |
| **Guillain-Barré**<br>**syndrome (GBS)**<br>**n = 23** | 9 / 14 | 51.7 y<br>(20 – 83) | 13.5 d<br>(1 – 65) | N. A. | PLEX (n = 10)<br>IVIG (n = 5) |
| **Myasthenia gravis**<br>**(MG)**<br>**n = 11** | 5 / 6 | 62.5 y<br>(30 – 89) | 23.2 d<br>(1 – 75) | Besinger<br>1.1<br>(0.2 –<br>2.3) | Mestinon (n = 9)<br>Prednisolon (n = 5)<br>Azathioprin (n = 2)<br>PLEX (n = 2)<br>IVIG (n = 2) |
| **Multiple Sclerosis**<br>**(MS)**<br>**n = 21** | 11 / 10 | 38.9 y<br>(22 – 65) | 16.2 d<br>(3 – 90) | EDSS<br>1.8<br>(0 – 4) | Methylprednisolone<br>( n = 4)<br>No other therapies before<br>blood draw |
| **Neuromyelitis**<br>**optica spectrum**<br>**disorder**<br>**(NMOSD)**<br>**n = 10** | 7 / 3 | 41.5 y<br>(21 – 82) | 2.3 y<br>(0.8- 5.8) | EDSS<br>1.3<br>(0 – 9) | Rituximab (n = 2)<br>Azathioprin (n = 1)<br>Glatirameracetat (n = 1)<br>PLEX (n = 3) |
| **Rheumatic**<br>**diseases**<br>**n = 11** | 6 / 5 | 50.7y<br>(18 – 74) | 50.7 y<br>(18 – 74) | N. A. | Methotrexat (n = 1)<br>Mycophenolat mofetil (n =<br>1)<br>Rituximab (n = 1)<br>Prednisolon (n = |

For vaccination studies, subjects received scheduled vaccinations following the German vaccination guidelines (STIKO). Altogether, 22 subjects received a vaccination against influenza (9 subjects Afluria 2014/2015, bioCSL; 13 subjects Afluria 2015/2016, bio CSL), 6 subjects received vaccination against tick borne encephalitis virus (FSME-Immun, Baxter,

### 3.2 Specimen handling, cell staining and sorting of peripheral blood B cell populations

Peripheral blood (25ml, EDTA blood) was collected from all patients during their routine diagnostic work-up. For vaccination experiments, blood was drawn from subjects before and at day-3, day-7, and day-14 after vaccination. In order to conduct several experiments after TBE vaccination, an additional time point for blood collection was day-9 for single subjects. Quantitative FACS analysis of B cell subtypes was performed within 6 hours after blood collection for all samples; for vaccination experiments we additionally obtained peripheral blood mononuclear cells (PBMCs, using a Ficoll gradient protocol, stored in liquid nitrogen) and serum used at a later time points for additional experiments.

For flow cytometric analyses, the following antibodies were used for all analyses: CD38 FITC (BD), IgD PE (Biozol), CD19 ECD (Beckman Coulter), CD3 PeCy7 (Beckman Coulter), CD45 VM (BD), CD27 APC (BD), CD20 APC Cy7 (BD). Accordingly, B cell subsets were defined by the following markers: naïve B cells CD19^+^CD20^+^CD27^−^CD38^+^IgD^+^, memory B cells CD19^+^CD20^+^CD27^+^CD38^+^, double negative B cells (DN B cells) CD19^+^CD20^low^CD27^−^IgD^−^, plasmablasts CD19^+^CD20^low^CD27^+^CD38^high^IgD^−^ (gating strategy is shown in suppl. Figure S2). Cell staining was visualized by additionally applying Image Stream for single samples.

For quantification of the different B cell subsets, immediate FACS analyses of fresh EDTA blood were performed on a CyAN ADP (Beckman Coulter) as described previously (34). For whole transcriptome and targeted transcriptome analyses, collected PBMCs from vaccinations were thawed at 37°C, washed in phosphate buffered saline (PBS) containing 2% FCS and incubated with the aforementioned antibodies (40min at 4°C). After another washing step, B cell subtypes were bulk sorted on a FACSAriaIII (BD bioscience) and collected in PBS directly followed by a RNA extraction step (Qiagen RNeasy Plus Micro Kit, manufacturer’s instructions, RNA stored at −80°C). For cloning of recombinant antibodies, single DN B cells and plasmablasts were sorted into 96 well plates on a FACSAriaIII (BD bioscience) and further processed as described below.

### 3.3 Whole transcriptome analyses

After cell sorting and RNA extraction, total RNA (about 200 pg) was amplified using the Ovation Pico WTA System V2 in combination with the Encore Biotin Module (Nugen). Amplified cDNA was hybridized on an Affymetrix Human Gene 2.0 ST array. Staining and scanning was done according to the Affymetrix expression protocol including minor modifications as suggested in the Encore Biotion protocol. Differential gene expression and statistical analyses were performed with an in-house R script (R v3.6.1) (35), mainly with the packages limma (v3.42.0), oligo (v1.50.0), and affycoretools (v1.58.0). The gene name annotation was performed with AnnotationDbi (v1.48.0) employing the Affymetrix hugene20 annotation data (v8.7.0). Genewise testing for differential expression was done employing the (paired) limma t-test and Benjamini-Hochberg (BH) multiple testing correction. Genes were considered differentially expressed which showed an adjusted p-value < 0.05 and log fold changes greater than 2. Heatmaps were generated with the pheatmap R package (v1.0.12). In one subject we could not access a representative transcriptome repertoire for the DN B cell population due to low RNA levels.

### 3.4 Targeted Ig transcriptome library preparation and sequencing

Next generation sequencing of Ig heavy chain VH transcripts was performed as described previously (6). Shortly, after cell sorting and RNA extraction (Qiagen RNeasy Plus Micro Kit, manufacturer’s instructions), cDNA synthesis was done using the Clontech SMARTer Ultra Low RNA Kit for Illumina sequencing according to the manufacturer’s instructions. We changed the second strand synthesis by additionally adding constant region primers for IgA, IgG, IgM and IgD to specifically pre-amplify immunoglobulin transcripts. After cDNA synthesis, a pool VH-family-specific (VH1-VH5) and isotype-specific (IgD, IgM, IgG, and IgA) primers were used to amplify VH-region sequences using polymerase chain reaction (PCR high fidelity, Roche); separate PCR reactions for each VH-family were performed to avoid cross-priming or primer competition. Ig constant region primers contained a sequence tag (“barcode”) to identify the cell population of origin (36). A primer sequence containing unique molecular identifiers (UMI) for subsequent Illumina MiSeq deep sequencing was included in the PCR primers, to control for sequence duplicates coming from the PCR amplification step. Amplified cDNA from the peripheral blood of each subject at each time point was then pooled and sequenced in a single run on an Illumina MiSeq Personal Sequencer, using 250bp paired-end sequencing.

On average, 184051 assembled heavy chain (VH) sequences (range: 77572–506890 sequences) were assessed on nucleotide level and further processed through the pRESTO bioinformatics pipeline to determine repertoires for each B cell population and time point (Suppl. Table S4). In consecutive steps, the total number of unique pRESTO pipeline VH sequences and clones were determined (Suppl. Table S4).

### 3.5 Targeted Ig transcriptome data analysis

The sequencing data was processed using the nf-core Bcellmagic pipeline (release 1.2.0), which is open source and available at http://github.com/nf-core/bcellmagic as part of the nf-core project (37). The pipeline employs the Immcantation toolset for processing of the repertoire sequencing data. The Illumina MiSeq high-throughput sequencing reads were quality-controlled using of FastQC. The pRESTO toolset (38) was used for processing the sequencing reads. Reads were filtered according to base quality (quality score threshold of 20), the forward and reverse reads were paired and a consensus sequence from reads with the same UMI barcodes was obtained, allowing a maximum mismatch error rate of 0.1 per read group. V(D)J sequences were only considered that had at least 2 representative sequences to build the consensus. Sequence copies were calculated as the number of identical sequences with different UMI barcodes.

VH variable-diversity-joining [V(D)J] germline segments were assigned by blasting the processed sequences to the IMGT database by igBLAST (39). Functional V(D)J sequences were assigned into clones based on (1) identical nucleic acid complementarity determining region-3 (CDR3) sequence length, (2) same VH variable gene segment and VH joining gene segment and (3) 88% identity of CDR3 nucleotide sequence. Clonal lineage reconstruction was performed with ChangeO (40) and by PHYLIP (for maximum parsimony linage construction) (41). Repertoire characterization and mutation profiling was performed by the use of Alakazam and SHazaM, respectively. Clonage tree graphs were exported in graphml format and loaded into a neo4j graph database (42), for automated lineage tree topology analysis. All tools were containerized in a singularity container distributed together with the analysis pipeline.

### 3.6 DN B cell single cell analysis, production of recombinant antibodies (rAb) and antibody testing

We performed single cell sorting with the aforementioned antibody panel in order to sort single double negative B cells; two PMBC samples (subject 1 and subject 5) obtained at day-9 after vaccination were sorted into double negative B cells (two 96 wells plates for each subject). Double negative heavy-(VH) and light-chain (VL) variable region sequences were recovered by RT-PCR and DNA sequencing (Eurofins, Munich, Germany) as described previously. Heavy- and light-chain sequences of 4 corresponding heavy- and light chains found within related clones (4 derived from double negative B cells) were selected, synthesized (Thermo Fisher Scientific Geneart, Regensburg, Germany) and introduced into expression vectors pIgG1Flag11 and pCEP4 (43,44); the vector for the heavy chain already contained the heavy chain constant region whereas the light chain was synthesized entirely and then introduced into the pCEP4 vector. Constructs were then co-transfected into HEK cells and antibodies produced commercially (EMP Genetech, Ingolstadt). Antibodies were purified (protein A columns) and run under reduced conditions on an SDS Gel obtaining heavy chain bands at around 50kDA and light chain bands at around 25kDA for 3 out of 4 antibodies (all derived from DN B cells). For one antibody, we just obtained a protein fragment of around 50kDA. Under un-reduced conditions, we identified a 160kDA band for 3 out of 4 antibodies; we concluded that these 3 antibodies were structurally intact.

In order to check the produced antibodies for antigen specificity, we performed specific enzyme immunoassays to measure IgG-antibodies against tick borne encephalitis in human serum (Immunozym FSME IgG, REF 7701010, PROGEN, Heidelberg, Germany). The ELISA tests were performed according to the manufacturer’s instructions. Shortly, wells of the ELISA test stripes are coated with inactivated tick-borne encephalitis virus and were then incubated with the recombinant antibodies or subject serum. Serum samples were diluted 1:100, antibodies were applied undiluted at a concentration between 0.1 mg/ml to 0.15 mg/ml. All 3 functional antibodies were tested as well as subjects’ serum samples at baseline and day-14 after vaccination. Standard curves showed a linear range and positive controls were within the recommended range. Results for recombinant antibodies were confirmed with a second ELISA test (Anti-FSME/TBE Virus ELISA “Vienna” (IgG) assay, EI 2661-9601-9 G, EUROIMMUN, Lübeck, Germany; application according to manufacturer’s instruction, dilutions identical first ELISA tests).

### 3.7 Statistics

Statistics for flow cytometry experiments were conducted using GraphPad Prism® Version 5.01. For cross sectional analysis of inflammatory neurological diseases the non-parametric Kruskal-Wallis test with multiple comparison correction (Dunn’s procedure) against control (NIND) was applied. For longitudinal analysis of vaccination experiments, repeated measurements ANOVA with Dunnett’s multiple comparison test between day-7 and baseline was applied. Values were considered statistically significant when p < 0.05. Indications: * p < 0.05, ** p < 0.01 and *** p < 0.001.

## 4 Results

### 4.1 Elevated peripheral blood DN B cells in active neuro-inflammatory diseases

We performed a cross sectional flow cytometric analysis of peripheral blood naïve, memory, DN B cells, and plasmablasts in patients with NMOSD, myasthenia gravis (MG), meningitis / encephalitis, Guillain-Barré syndrome (GBS), MS, non-inflammatory neurological diseases (NIND) and rheumatic diseases. The overall B cell pool, calculated as the percentage of CD19^+^ cells of all mononuclear cells (CD45^+^), did not show significant differences between patient groups (mean 11% for each patient group, range 8-13%). However, differences were observed for the distribution of B cell subtypes including naïve B cells, memory B cells, DN B cells and plasmablasts (calculated as the percentage of all CD19^+^ B cells). Higher DN B cells were detectable in patients with MG, meningitis / encephalitis, GBS rheumatic diseases, and NMOSD (Figure 1A). Peripheral plasmablasts showed significantly higher values in patients with GBS, meningitis / encephalitis, MS and NMOSD when compared to NIND (Figure 1A). Besides a decreased percentage of memory B cells in rheumatic diseases, no significant changes were detectable for memory or naive B cells (Figure 1A).

**Figure 1:**
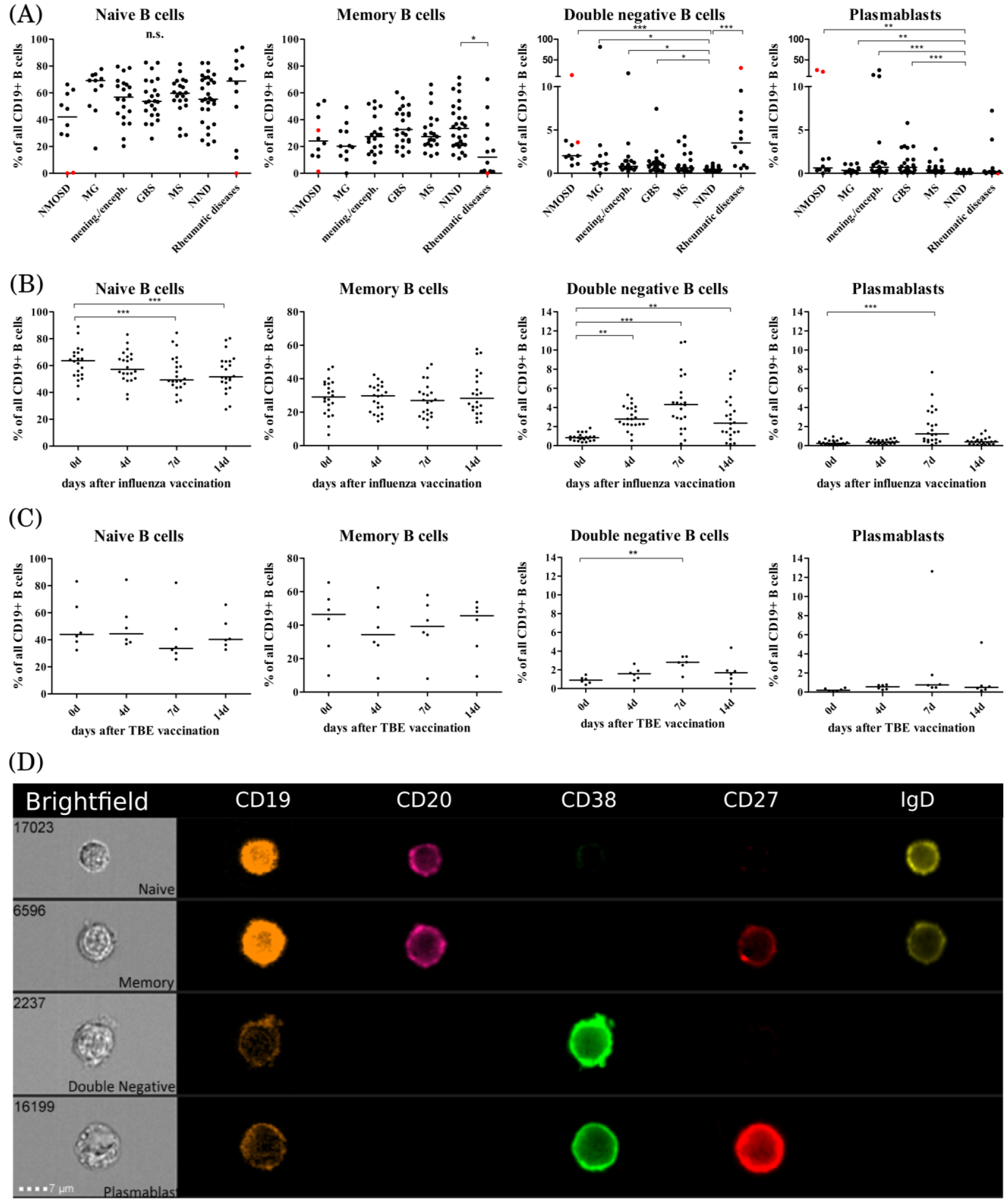
Distribution of B cell subsets in the peripheral blood of patients with neuro-inflammatory disorders or following vaccination with influenza or tick borne encephalitis virus. B cell subsets are presented as a percentage of total CD19^+^ B cells and defined as follows: naïve B cells CD19^+^CD20^+^CD27^−^CD38^+^IgD^+^, memory B cells CD19^+^CD20^+^CD27^+^CD38^+^, double negative B cells (DN B cells) CD19^+^CD20^low^CD27^−^IgD^−^, plasmablasts CD19^+^CD20^low^CD27^+^CD38^high^IgD^−^. (**A**) Peripheral blood B cell subsets were quantified from patients with neuromyelitis optica spectrum disorder (NMOSD), myasthenia gravis, meningitis / encephalitis (mening./enceph.), Guillain-Barré syndrome (GBS), multiple sclerosis (MS), non-inflammatory neurological diseases (NIND) and patients with rheumatic diseases. Red dots highlight samples obtained from rituximab-treated patients. Clinical groups were compared using a Kruskal-Wallis test with correction for multiple comparisons. Peripheral blood B cell subsets following vaccination against (**B**) influenza virus (n = 22) or (**C**) tick-borne encephalitis virus (n = 6). Statistical measurements were performed using a repeated measures ANOVA with Dunnett’s post-hoc correction. (**D**) Staining of different B cell populations was visualized by ImageStream. Lines in the graphs indicate median values and asterisks describe significance values as follows: * p < 0.05, ** p < 0.01, *** p < 0.001

Statistically significant differences were not noticeable between disease subgroups. This may be due to heterogeneity in disease duration (e. g. GBS), disease severity, time of last relapse, and immunomodulatory therapies (patient characteristics Table 1). We did not find a correlation between the percentage of DN B cells and age (Spearman test, p = 0.3).

### 4.2 Peripheral blood DN B cells are upregulated after vaccination

In order to further study and characterize DN B cells, we monitored the specific PB B cell response after vaccination against influenza virus (n = 22) or tick-borne encephalitis virus (n = 6) in healthy subjects (subject characteristics and conducted experiments are summarized in Table 2). The distribution of naïve, memory, DN B cells and plasmablasts was calculated as percentage of all B cells. We observed an increase in circulating DN B cells day-4, -7 and -14 after vaccination against influenza virus (Figure 1B), maximum at day-7. In addition, plasmablasts were significantly upregulated on day-7 whereas a significant percentage decrease of naive B cells was observed on days-7 and -14. A similar increase of DN B cells at day-7 was detectable after vaccination against tick born encephalitis virus (TBE) (Figure 1C). To further confirm the validity of our FACS protocol, we visualized individual B cell populations using live imaging (ImageStream; Figure 1D).

**Table 2:**
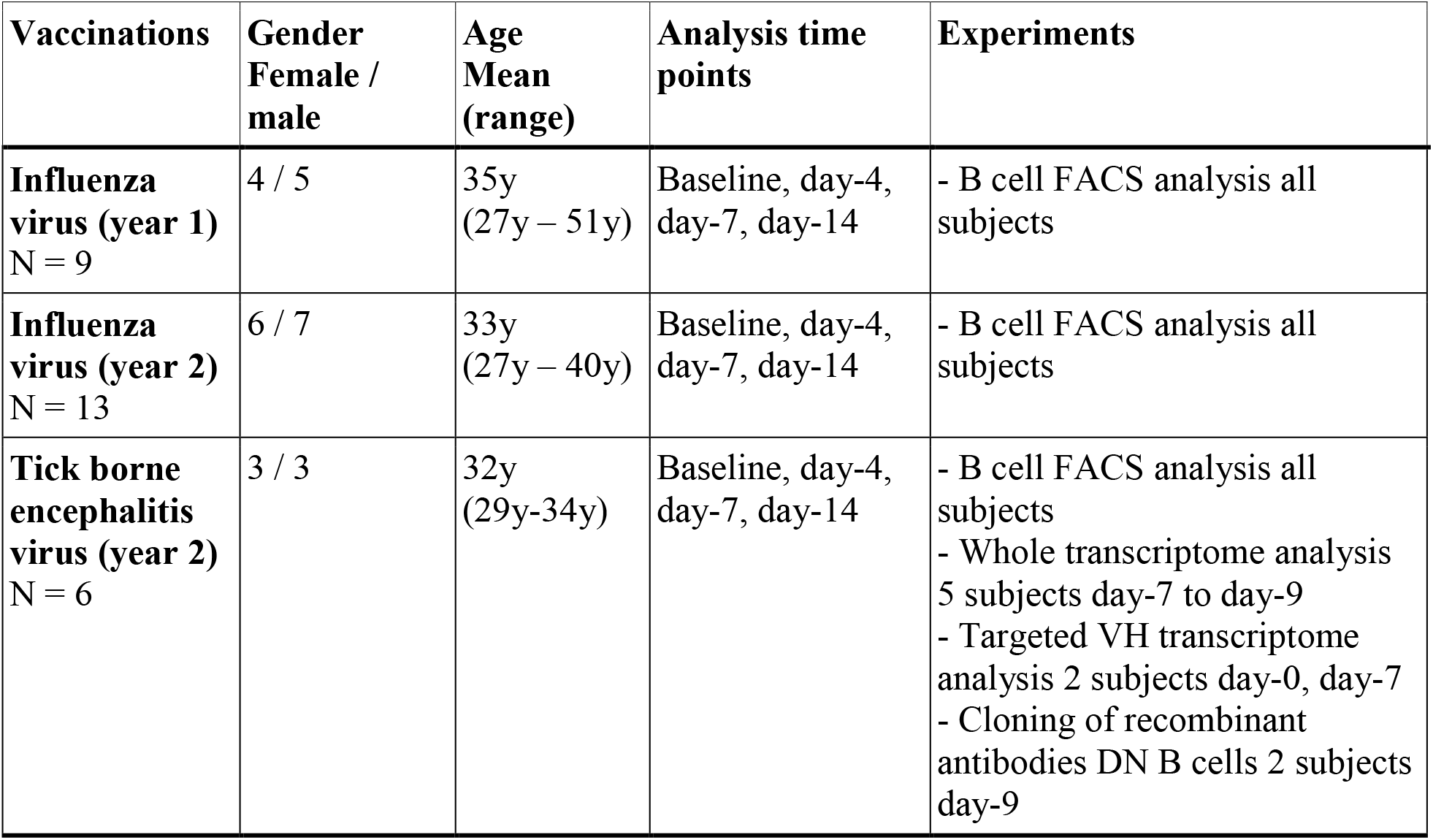
Characteristics of healthy subjects receiving vaccinations against influence and tick borne encephalitis virus and conducted experiments.

| **Vaccinations** | **Gender**<br>**Female /**<br>**male** | **Age**<br>**Mean**<br>**(range)** | **Analysis time**<br>**points** | **Experiments** |
| --- | --- | --- | --- | --- |
| **Influenza**<br>**virus (year 1)**<br>N = 9 | 4 / 5 | 35y<br>(27y – 51y) | Baseline, day-4,<br>day-7, day-14 | - B cell FACS analysis all<br>subjects |
| **Influenza**<br>**virus (year 2)**<br>N = 13 | 6 / 7 | 33y<br>(27y – 40y) | Baseline, day-4,<br>day-7, day-14 | - B cell FACS analysis all<br>subjects |
| **Tick borne**<br>**encephalitis**<br>**virus (year 2)**<br>N = 6 | 3 / 3 | 32y<br>(29y-34y) | Baseline, day-4,<br>day-7, day-14 | - B cell FACS analysis all<br>subjects<br>- Whole transcriptome analysis<br>5 subjects day-7 to day-9<br>- Targeted VH transcriptome<br>analysis 2 subjects day-0, day-7<br>- Cloning of recombinant<br>antibodies DN B cells 2 subjects<br>day-9 |

### 4.3 DN B cells do not uniquely cluster by gene expression

To characterize the gene expression profile of the different B cell subtypes, we performed a whole transcriptome analysis of naïve, memory, DN B cells and plasmablasts from five subjects following vaccination against tick borne encephalitis virus.

Principal component analysis (PCA) showed clustering of naïve and memory B cells, whereas plasmablasts formed a separate cluster that can be distinguished according to values of the first principal component (PC1) (Figure 2A). DN B cells, on the contrary, did not show clear clustering according to their gene expression profile. This cell type clustered with naïve and memory B cells in three subjects, and together with plasmablasts in one subject (Figure 2A). We could not access a representative transcriptome analysis for the DN B cell population for subject S5 due to low RNA levels.

**Figure 2:**
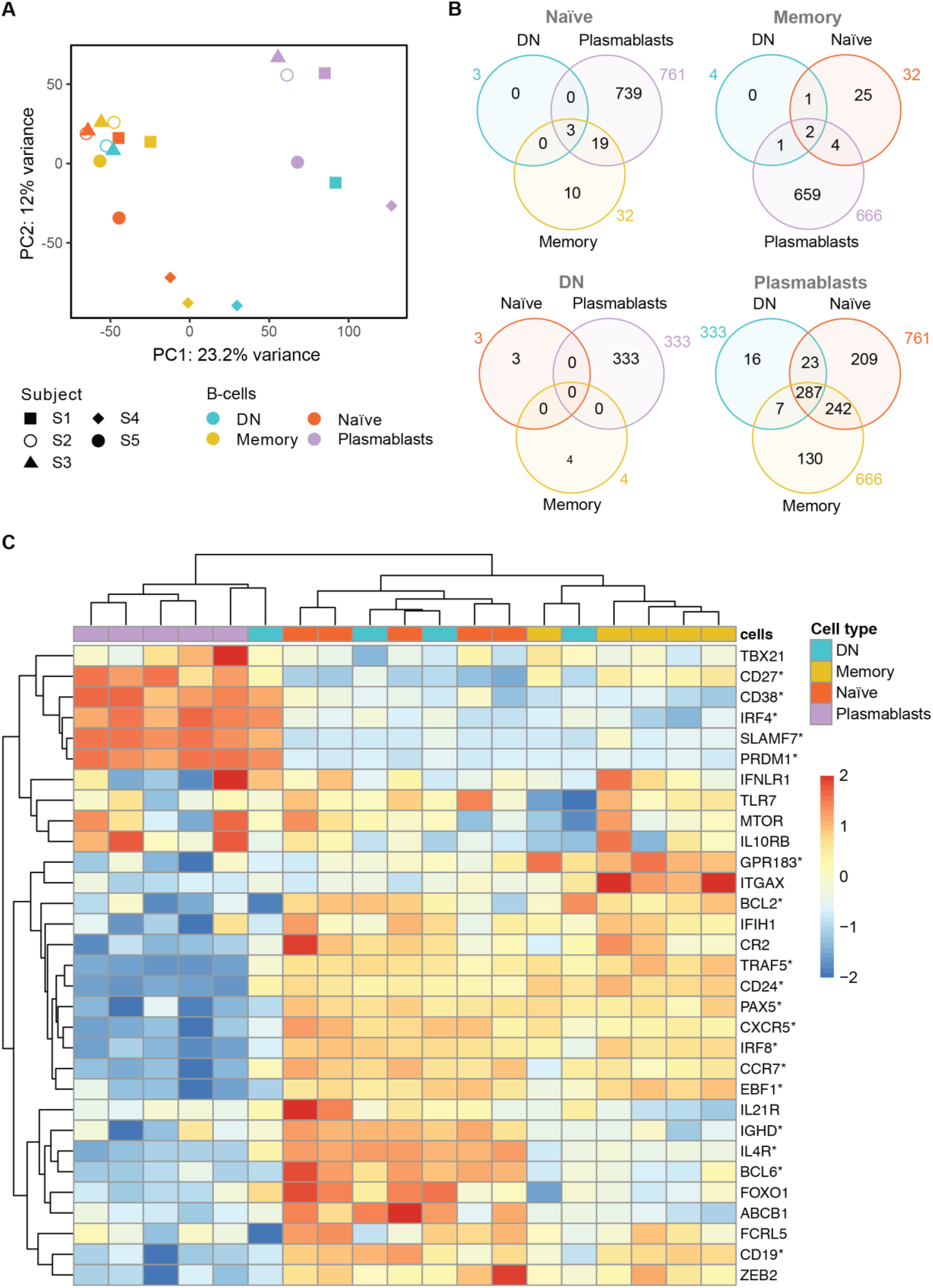
Whole transcriptome analysis of the different B cell populations after TBE vaccination (**A**) Principal component analysis of whole transcriptome data with clustering of different B cell subsets by subjects (S1-S5). Plasmablasts (purple) cluster separately from memory (yellow) and naive (red) B cell subsets whereas DN B cells (blue) show an inconsistent clustering with naïve / memory B cells, plasmablasts or between the latter population. Inter-individual differences were noticeable especially for subject 4 (S4). (**B**) Venn Diagrams of differentially expressed genes (DEG, p<0.05; fold-change >2x) of shared between the B cell types for each baseline comparator population (indicated above in gray). The number of DEG was limited (≤ 32) between naïve, memory and DN B cells; plasmablasts showed > 300 DEG when compared to any of the other B cell subtypes with DN B cells showing the lowest number of subset specific DEG. (**C**) Heatmap displaying the normalized expression values of selected genes of interest across the different B-cell types. Significantly DEG are marked by asterisks as detailed in Supplementary Table S1. The color scale shows the normalized gene expression values scaled between −2 and 2 for each gene (row). Hierarchical clustering dendrogram of B-cell subsets according to their gene subset expression profiles (top) and clustering dendrogram of the genes according to their expression patterns in the different samples (left) are shown. A consistent grouping of plasmablasts, naïve B cells and memory B cells can be found with DN B cells showing an inconsistent grouping with each of the other B cell subsets. Abbreviations: subject 1-5 = S1-S5, DN = DN B cells.

Next, we analyzed differentially expressed genes (DEG) between the four B cell populations using each B cell subset transcriptome to define the baseline comparator for the other three subsets (Figure 2B). When comparing the gene expression among naïve, memory and DN B cell populations, only a limited number of DEG was found (≤ 32 genes) for any of the comparisons. In contrast, plasmablasts showed ≥333 DEG when compared to each of the other B cell subtypes. Using plasmablasts as the baseline, DN B cells showed the lowest number of subsets specific DEG (16) followed by memory B cells (130) and naïve B cells (209) beside multiple DEG shared between the latter populations.

Specific analyses of annotated, differentially expressed transcripts that were only found between DN B cells and the other populations were further conducted. We identified IGH (Ig heavy locus) and IGHA1 (Ig heavy chain constant region alpha1) as DEG between DN and naïve B cells with both transcripts showing a significantly higher expression in DN B cells (Suppl. Fig. S2 A, DEG between memory and naïve B cells additionally shown in Fig. S2 B). Between DN and memory B cells, TLR6, IGHV4-31 and TFEC showed a significantly different expression (Suppl. Fig. S2 C). The TLR6 (toll like receptor 6) gene showed a significantly higher expression in DN B cells which might be associated with the specific response against viruses in combination with the TLR2 (45). The upregulation of VH4 gene segments in DN B cells (especially IGHV4-34) has previously been reported in SLE (2). Transcriptional factor EC (TFEC) is associated with an immunoglobulin heavy-chain gene enhancer (46) and was upregulated in memory B cells. Detailed analysis of annotated DEG between DN B cells and plasmablasts revealed a significant higher expression of IGKV2-29 in plasmablasts (data not shown). Furthermore, DN B cell specific DEG contained RNUF5F-1, RARRES3, MBNL2, MRPL12, NDUFA7 but no obvious B cell associated functions could be assigned to these genes.

In addition, we examined the differential expression of 31 genes of interest that have previously been associated with B cell development and DN B cells (Figure 2C, Suppl. Table S1). Hierarchical clustering grouped plasmablasts, naïve B cells and memory B cells, consistent to the PCA analysis. In line with the principal component analysis, DN B cell subsets again showed inconsistent mRNA expression patterns with three DN B cell populations clustering with naïve and memory B cells, and one with plasmablasts. Also, CD27 and CD38 showed a significantly higher gene expression in plasmablasts than in DN and naïve B cells. CD24, an early B cell marker (47), was expressed at significantly lower levels in plasmablasts than in DN, memory and naïve B cells. Further sub-classification of DN B cells by the chemokine receptor CXCR5 and integrin CD11c (ITGAX) into DN1 (CXCR5+, CD11c) and DN2 (CXCR5-, CD11c++) has recently been suggested by Sanz and colleagues (1). We observed a significantly lower expression of CXCR5 transcripts in plasmablasts when compared to the other cell types; no significant differences were observed for CD11c transcripts between the different subpopulations. When looking at DN B cells on a subject level, we found a down-regulation of CXCR5 transcripts similar to plasmablasts in one subject and a higher expression CD11c transcripts in DN than in naïve B cells but differences did not reach significance. Concerning cell to cell signaling and cell development, we found a higher expression of IL4R transcripts in DN and naïve B cells when compared to plasmablasts. Recently it could be shown, that IL4 together with IL21 and INFγ control CD11c and TBET (TBX21) expression in B cells (3). TNF receptor associated factor (TRAF5) transcripts that mediate TNF induced cell activation (3) showed a lower expression in plasmablasts when compared to the other cell populations; also DN B cells showed low expression levels on a subject level which is in line with the literature. IRF4 as a transcription factor for plasma cell differentiation was expressed in plasmablasts at higher levels than in naïve and memory B cells but not DN B cells (3). The transcripts PRDM1 (Blimp1) and SLAMF7 are also associated with plasma cell differentiation and showed a higher expression in plasmablasts when compared to the other populations, however, DN B cells in one subject also showed high expression of both transcripts similar to plasmablast levels (further details are shown in Suppl. Table S1).

### 4.4 Targeted immunoglobulin repertoire analyses show clonal expansion of DN B cells

We performed targeted heavy chain (VH) transcriptome sequencing in two subjects (S1 and S5, part of the aforementioned analysis) at baseline and 7 days after vaccination against TBE. Basic VH transcriptome parameters were studied including VH family usage, Ig subclass distribution, and mutational count (Suppl. Table S2). The VH family distribution of naive B cell repertoires at baseline approximated the expected germline frequency (48), however, at day-7 after vaccination, naïve B cells showed a higher diversity. Concerning the other B cell subsets, DN also showed a VH family usage close to germline frequencies whereas memory B cells and especially plasmablasts showed a heterogeneous VH usage. Regarding Ig isotypes, the frequency of IgD was lower in the naïve population following vaccination at day-7 with IgM being overly expressed; no obvious trends were observed for the other populations. The mean number of mutations was lowest in the naïve populations indicating representative sequencing of VH transcripts for the different B cell subsets. There was a tendency for an increased mean mutational count after vaccination, especially for DN B cells (further details Suppl. Table S2).

When analyzing the clonal diversification of the BCR repertoire for the different B cell populations, we found a significantly lower diversification for all B cell subsets after vaccination except for DN B cells at day-7 (Figure 3 A, Suppl. Fig. S3). The clonal diversification index measures the unevenness of the number of unique BCR sequences per clone (49), and an increased diversification after vaccination indicates clonal expansion. Further evidence for a clonal expansion of the DN B cell subset derived from the percentage distribution of clones with 10 or more sequences (Figure 3B, Suppl. Fig. S3 and S4). Clonal abundance significantly increased in the DN B cell population as well as in the memory and plasmablast subsets but decreased or remained unchanged for naïve B cells. We next analyzed clonally related VH sequences within and between the different B cell subsets at baseline and day-7 after vaccination (Figure 3C). The majority of clonally related VH sequences were found within each B cell population itself. The percentage overlap between the different populations increased after vaccination from 2.1% to 23.8% in subject 1 and 5.3% to 10.7% subject 5 after vaccination. In subject 1, the percentage overlap of DN B cells to all other B cells subsets increased, including naïve B cells. In addition, naïve B cells also showed direct clonally related sequences to plasmablasts. An increase in overlapping sequences between DN B cells and memory B cells and plasmablasts was also observed in subject 5. However, the overlaps between naïve and other B cell subtypes remained scarce at day-7 in subject 5 (Figure 3C and Suppl. Fig. S5).

**Figure 3:**
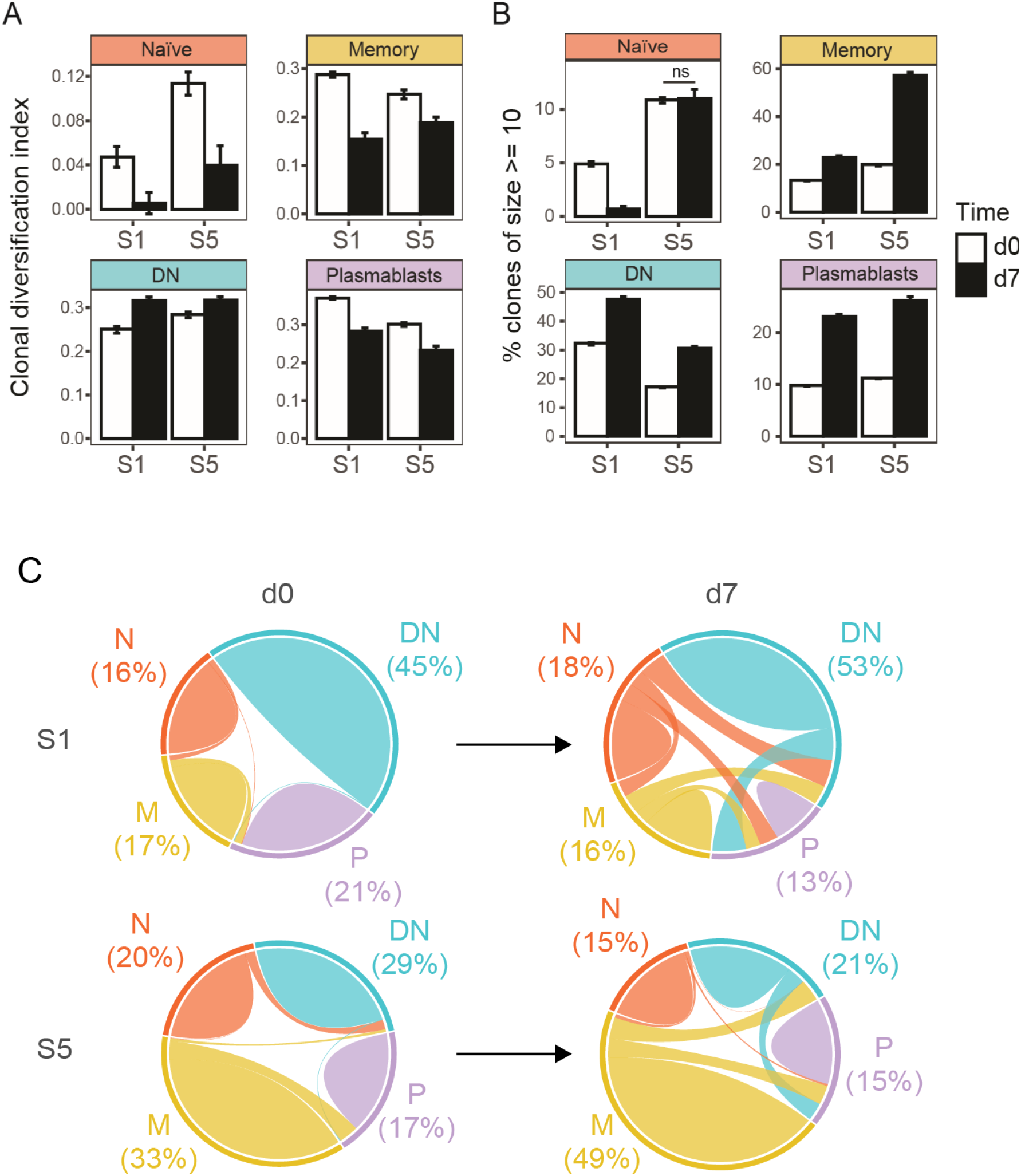
Targeted immunoglobulin VH transcriptome analysis in subjects 1 (S1) and 5 (S5) before (d0) and after 7 days (d7) of TBE vaccination. (**A)** Clonal diversification amongst clonal populations at baseline and day-7 after vaccination is shown for the different B cell subsets. (**B)** Percentage of large clonal populations (≥ 10 unique sequences) in circulating B cell subsets. Clonal diversification index and the percentage of large clones were calculated using a sample of 611 clones collected in accordance with their abundance in the repertoire (n=1000 bootstrap iterations). The error bars indicate standard deviation. All pair-wise comparisons were significant (p < 0.001, t-test) between the bootstrap samples unless otherwise indicated (ns: not significant). (**C)** Overlap analyses (weighted by the number of unique sequences) of clonally related sequences within and between the different B cell populations. Numbers indicate the percentage of sequences from each B cell subtype for each patient and time point. Abbreviations: N: naïve, M: memory, DN: DN B cells, P: plasmablasts.

Directed maturation trees were constructed by aligning clonally related VH sequences to their most homologous germline. The succession of B cell subtypes defined by the patterns of somatic hypermutations was then used to predict the most likely direction of B cell maturation after vaccination (e.g. DN B cells followed by plasmablasts, Figure 4). At baseline, a very limited number of maturation trees containing several B cell subtypes (subject 1: 0 trees, subject 5: 7 trees) was observed whereas multiple trees were found (subject 1: 74 trees, subject 5: 55 trees) after vaccination. In detail, maturation trees containing naïve B cells (n = 31), DN B cells (n = 29) and memory B cells (n = 7) were directly followed by plasmablasts in subject 1. Memory B cells (N = 31) and DN B cells (n = 7) followed by plasmablasts as well as memory B cells (n = 7) followed by DN B cells mainly contributed to directed maturation trees in subject 5.

**Figure 4:**
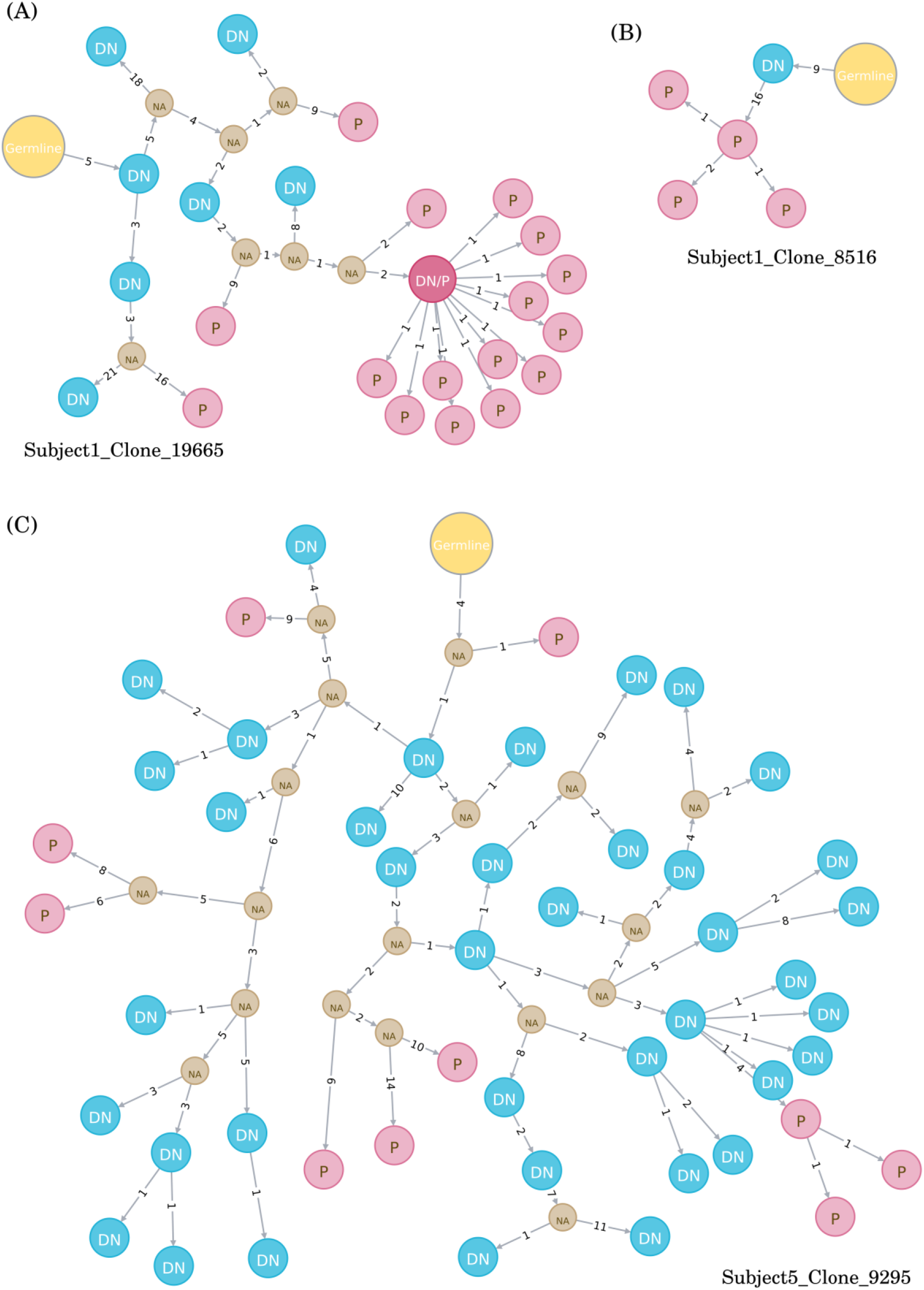
Examples of directed maturation trees (all at timepoint day-7). Clonally related VH sequences were aligned to their most homologous germline and maturation pathways and the succession of B cell subtypes defined by the patterns of somatic hypermutations. DN B cells (blue circles) often precede plasmablasts (red cicle) that show more somatic hypermutations. Abbreviations: NA = unknown intermediates, DN = DN B cell, P = plasmablast

### 4.5 DN B cell derived, recombinant antibodies show specificity against TBE

We generated 3 recombinant antibodies (2 from S1; 1 from S5) from over-represented clonally related Ig sequences expressed by single DN B cells at day 9 after vaccination against tick borne encephalitis. The specificity of these antibodies was tested by commercially available ELISA tests that are used to evaluate serum IgG titers after vaccination. When measuring IgG serum titers in both subjects, we observed negative titers in S1, borderline titers in S5 at baseline and sufficient anti-TBE-IgG titers at day-14 after vaccination for both subjects (Table S3). One recombinant antibody from S1 (F4) showed a mean value of 138VIEU/ml (titers considered positive when > 126 VIEU/ml) whereas the second antibody (C3) showed mean values of 90 VIEU/ml (borderline 63 – 126 VIEU/ml) indicating that both DN B cells produced TBE specific antibodies. The third antibody from S5 (D10) was below the detection limit of the ELISA test (Table S3). Results were confirmed using a second ELISA Kit.

## 5 Discussion

DN B cells have been suggested to contribute to the pathophysiology of several autoimmune disorders including (NMOSD) (*6*) and SLE (2,10–12) but also infectious diseases like HIV (17–19), sepsis (20) and active malaria (21,22). However, little is known about the origin and function of DN B cells and the involvement in other inflammatory neurological diseases. In this study we show, that elevated peripheral blood (PB) DN B cells are observed in a variety of autoimmune and infectious neurological disorders and are robustly induced by vaccination. Further characterization of DN B cells after TBE vaccination by whole transcriptome analyses revealed a transcriptional program that overlaps with naïve B cells, memory cells, and plasmablasts. Targeted Ig transcriptome analyses in combination with the production of recombinant antibodies from DN B cells suggest an antigen driven maturation process upon certain immunologic stimuli.

In line with previous studies (6,11), we observed elevated values of peripheral blood DN B cells in patients NMOSD and rheumatic diseases. In addition, we found an increased fraction of DN B cells in patients with acute MG and GBS providing indirect evidence for a possible involvement of DN B cells in several autoantibody associated diseases. Under conditions of an acute infection in patients with meningitis / encephalitis, increased numbers of peripheral blood DN B cells were detectable similar to patients with acute sepsis (20–22). Similar to a recent report, only a subset of MS patients showed an elevated fraction of DN B cells (7). Although an elevated fraction of DN B cells has been reported in the elderly (>75y) (4,5), we did not observe a correlation between DN B cells and age in our patient cohorts. Following vaccination against influenza or TBE, we observed a robust expansion of DN B cells between day-4 and day-14, maximal at day-7. The increased fraction of DN B cells was accompanied by an expansion of plasmablasts at day-7 which is in line with the literature (50,51). Although detailed information is not available, a recent study also described the induction of DN B cells after vaccination with influenza virus (52). In summary, we found an increased fraction of peripheral blood DN B cells in autoimmune, infectious, and immunized patients, suggesting that DN B cells play an important role in both pathologic and protective, antigen-targeted immune responses.

A variety of functional roles have been proposed for DN B cells: exhausted memory B cells, transient effector B cells, or an atypical memory-like B cell population (32,33). Recent studies on DN B cells in SLE provide compelling evidence that a subset of DN B cells (DN2) represent a primed precursor population for antibody secreting cells (ASC) (1–3). Our detailed analyses of DN B cells after vaccination against TBE containing whole transcriptome analysis, targeted Ig VH repertoire sequencing and cloning of recombinant antibodies provides further evidence that DN B cells are a transient precursor B cell population undergoing differentiation into antibody secreting cells. Whole transcriptome analyses showed a heterogeneous transcriptome in DN B cells that showed multiple overlaps with naïve B cells, memory B cells and plasmablasts. Principal component analysis showed that DN B cells demonstrate a rapidly changing gene expression profile between naïve, memory and plasmablast subsets that did not coalesce into an independent cluster. While plasmablasts show a distinct gene expression profile in comparison to the other populations, the number of identified DEG was lower between plasmablasts and DN B cells. Indeed DN B cells expressed transcription factors like IRF4 which are induced during plasma cell differentiation (*3*). A direct clonal relation between DN B cells and plasmablasts was also observed by targeted VH transcriptome analyses. Following vaccination, DN B cell clones were observed to link directly to plasmablasts by lineage trees (Figure 4). Furthermore, recombinant antibodies generated from DN B cells recovered from TBE-vaccinated subjects bound to TBE-infected cell lysate by ELISA. In combination, our data indicates that DN B cells provide a distinct pathway for the generation of antibody producing plasmablasts in response to antigenic stimulation.

Sanz and colleagues recently provided evidence that DN2 B cells develop from the naïve B cell pool through an extra-follicular pathway (1–3). In contrast, DN1 B cells (CXCR5^+^, CD21^+^, CD11c^−^) comprised an activated subset of memory B cells arising through follicular development (3). Although our FACS approach did not allow us to differentiate DN B cells into DN1 and DN2 subsets, VH transcriptome analysis suggests a similar distinction amongst our post-vaccination DN B cells. Following vaccination, we observed an activation of the naïve B cell pool with a loss of IgD expression and greater bias in VH germline usage. The transcriptomes of naïve and DN B cells demonstrated few DEGs with elevated expression of IGH and IGHA1 in DN B cells indicating the onset of Ig class switching. Indeed, DN B cells were mostly class-switched (30 - 44% IgA-expressing) and showed an increased frequency of somatic mutations and large clonally-expanded populations. While toll-like receptors (TLR) 7 and 9 have recently been associated with the differentiation of B cells into DN B cells in SLE (3,53), we observed a high expression of TLR6 in our DN B cell subset after TBE vaccination. The TLR6 together with TLR2 has also been associated with virus recognition, which further argues for a TLR mediated activation in the context of DN B cell maturation (45). Further indirect evidence for extra-follicular DN maturation derives from the clonal connectivity between our different B cell subtypes. When looking at the specific clonal relations between the B cell populations after TBE vaccination, we found several clones between naïve B cells and plasmablasts / DN B cells as well as DN B cells and plasmablasts whereas the connections between memory B cells and the other populations remained scarce in subject 1. These clonal interactions between the different populations could point towards the aforementioned extra-follicular pathway with activated naïve B cells followed by DN B cells that develop into plasmablasts. However, we observed distinct differences in the clonal interaction between the different populations in subject 5. Most clonal interactions were found between memory B cells, DN B cells and plasmablasts whereas the naïve B cell population only showed limited clonal overlap with the other B cell subsets. However, subject 5 already showed intermediated anti-TBE-IgG titers at baseline, so that the clonal interaction could possibly resemble a reactivation of the B cell memory compound followed by DN B cell and plasmablast generation. Regardless of potentially different maturation pathways, both subjects developed a sufficient serum antibody response evaluated by a specific ELISA test. Altogether, the different clonal connectivity could be explained by the DN1 and DN2 concept but further experiments are necessary to fully address B cell maturation following vaccination.

The characterization of DN B cells in our vaccination model has indirect implications for the interpretation of elevated DN B cells in neurological diseases. We recently showed that the fraction of PB DN B cells is elevated during active NMOSD and contains AQP4 positive B cell clones (6). Similar to these results, DN B cells have been observed to express disease associated antibodies in SLE (3). Although we did not examine whether DN B cells in our patients with myasthenia gravis, Guillain-Barré syndrome or meningitis / encephalitis are associated with a specific, disease relevant antibody response, the acute and / or relapsing clinical course of these disorders suggests that DN B cells may represent a source of disease-relevant immunoglobulins. The interval between proceeding infections and the onset of GBS or relapse in MG has been reported to vary between 1 to 3 weeks (54,55), in addition GBS sometimes occurs after vaccinations (54). Given that the B cell / antibody response after vaccinations show a similar time frame, a potential involvement of DN B cells in the production of autoantibodies in MG and GBS seems plausible but needs further confirmation in future studies.

Some limitations of our study should be noted. We included CD20 in our FACS panel and followed a gating strategy (6) using CD20 for the definition of double negative B cells. We used this surface marker, since CD20 B cell depleting therapies are increasingly applied to autoimmune diseases. However, this gating strategy lead to a slightly different definition of DN B cells which makes it difficult to directly compare our flow cytometric results to other studies, although the DN B cell population has also been reported to be CD20 negative applying different gating strategies (1). Our patient cohort showed a certain degree of heterogeneity with respect to disease activity, duration and treatment which limits comparisons. For this reason, we characterized DN B cells after a defined immunologic stimulus in a vaccination model, similar to recent analyses of acute SLE patients (3). Methodological limitations included low patient numbers, low RNA levels for whole transcriptome analysis, and a limited sampling of targeted Ig transcriptome repertoires. Nevertheless, the results provide proof of concept for the induction of DN B cells post-vaccination, provide a robust characterization at the transcriptome level, and reveal targeted antigen specificity through recombinant antibody technology.

In summary, our study shows that DN B cells are involved in a variety of active neuro-inflammatory diseases as well as vaccinations thus playing a role in protective as well as pathogenic immune responses. Post-vaccination DN B cells comprised a transient population that developed into specific antibody secreting plasmablasts. Future studies with a sub-differentiation of DN B cells into the DN1 and DN2 phenotype are necessary to fully understand follicular and extra-follicular B cell maturation pathways and the exact role of DN B cells in specific neuro-inflammatory disease entities and during vaccinations.

## Author Contributions

**CR** collected and analyzed the data, did statistical analysis, did literature research, wrote the manuscript, critically reviewed and revised the manuscript; **GG** analyzed the data, performed statistical analysis, wrote the manuscript and critically revised the manuscript; **GL** analyzed the data, did statistical analysis (consulting statistician) and critically reviewed the manuscript for important intellectual content; **SH** curated the data for analysis; **MK** analyzed the data and critically reviewed the manuscript for important intellectual content; **ZH** performed and analyzed FACS analysis and critically reviewed the manuscript for important intellectual content; **MI** collected the microarray data and critically reviewed the manuscript for important intellectual content; **JB** supervised the microarray analysis and critically reviewed the manuscript for important intellectual content; **UZ** critically reviewed the manuscript for important intellectual content; **SN** analyzed the data (consulting statistician and senior bioinformatician) and critically reviewed the manuscript for important intellectual content; **GPO** critically reviewed the manuscript for important intellectual content; **JLB** analyzed the data and critically reviewed the manuscript for important intellectual content; **BH** critically reviewed the manuscript for important intellectual content; **MCK** designed and supervised the study, collected and analyzed the data, wrote the manuscript and critically revised the manuscript.

## Acknowledgments

We thank S. Poeschel and S. Autenrieth, FACS core facility Berg, University of Tübingen for expert assistance in acquisition of ImageStream flow cytometry. We Thank K. Thürmel, Department of Nephrology, Technische Universität München, Munich, Germany for helping with sample collection for patients with rheumatic diseases and F. Welser (EMP Genetech) for assistance with antibody production.

## Supplementary Material

Supplementary Figure S1: Gating strategy

Supplementary Figure S2: Normalized gene expression for DEG in 3 different contrasts

Supplementary Figure S3: Clonal abundance and clonal diversity

Supplementary Figure S4: Percentage of clones in the repertoire ≥C

Supplementary Figure S5: Clonal overlaps before and after vaccination

Supplementary Table S1: Significance of up/down-regulation of genes of interest

Supplementary Table S2: Base repertoire characteristics of targeted Ig repertoire analysis

Supplementary Table S3: Tick borne encephalitis ELISA of recombinant antibodies and serum

Supplementary Table S4: Cell counts, number of VH sequences and number of clones

