## Supplementary Figures and Tables for "Specific induction of double negative B cells during protective and pathogenic immune responses"

### Supplementary Materials:

#### Supplementary Figure S1: Gating Strategy

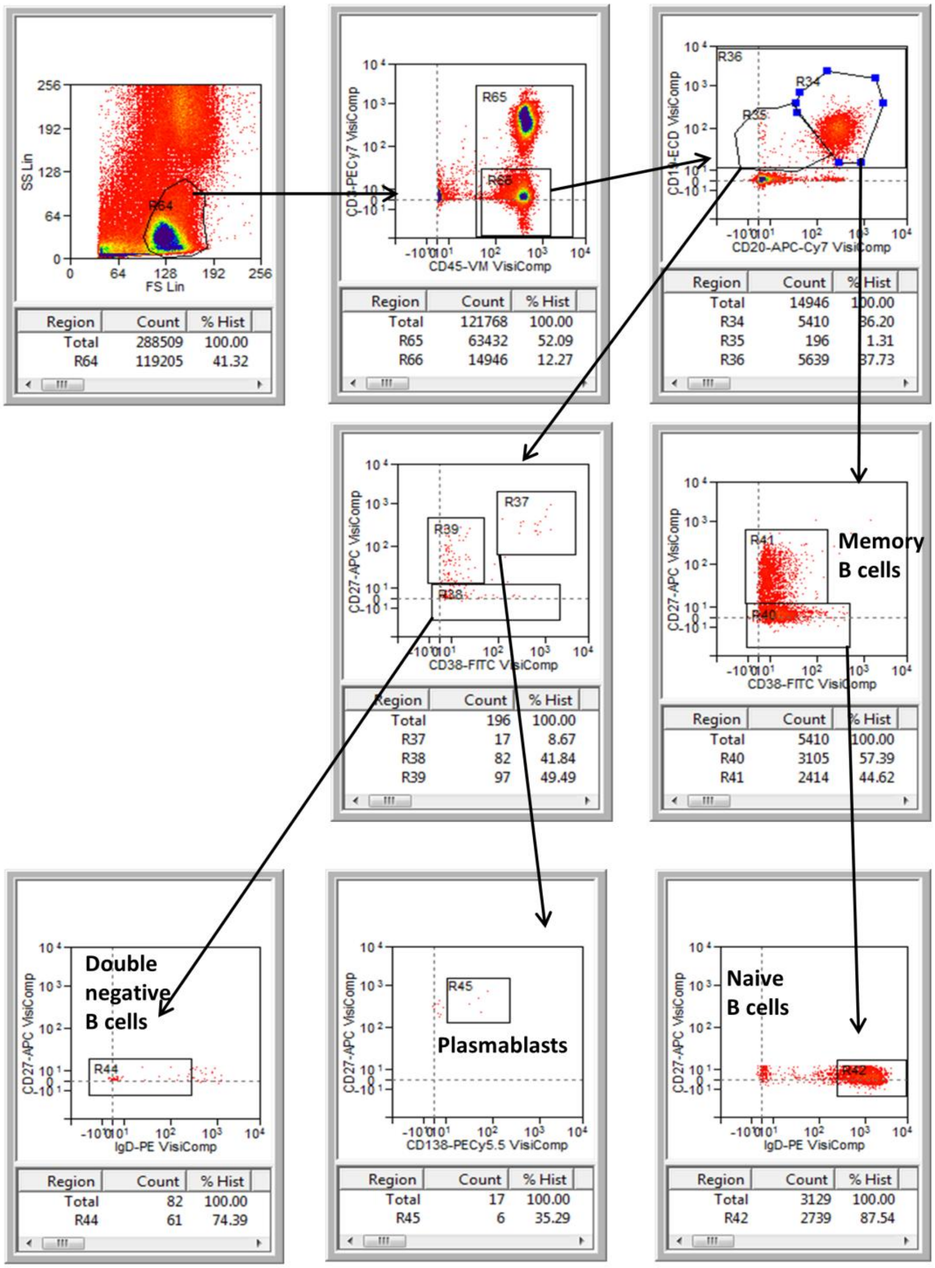

**Supplementary Figure S2**

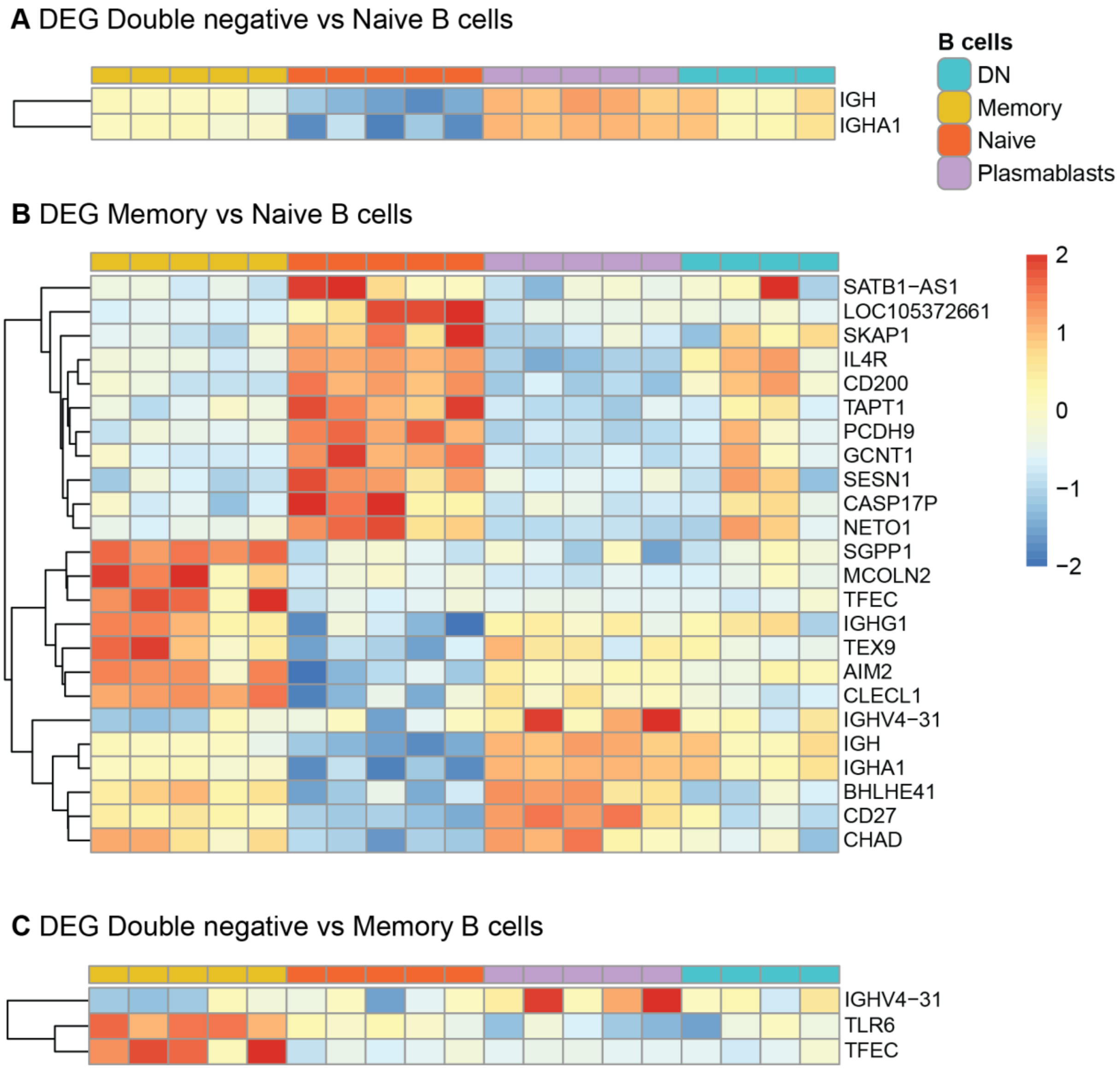


**Fig. S2:** Normalized gene expression for DEG in 3 different contrasts: (A) DN B-cells vs Naïve B-cells, (B) Memory vs Naïve B-cells, and (C) DN vs Memory B-cells. The hierarchical clustering dendrogram of the genes according to their expression patterns in the different samples is shown at the left. The color scale shows the normalized gene expression values scaled between -2 and 2 for each gene (row).

**Supplementary Figure S3**

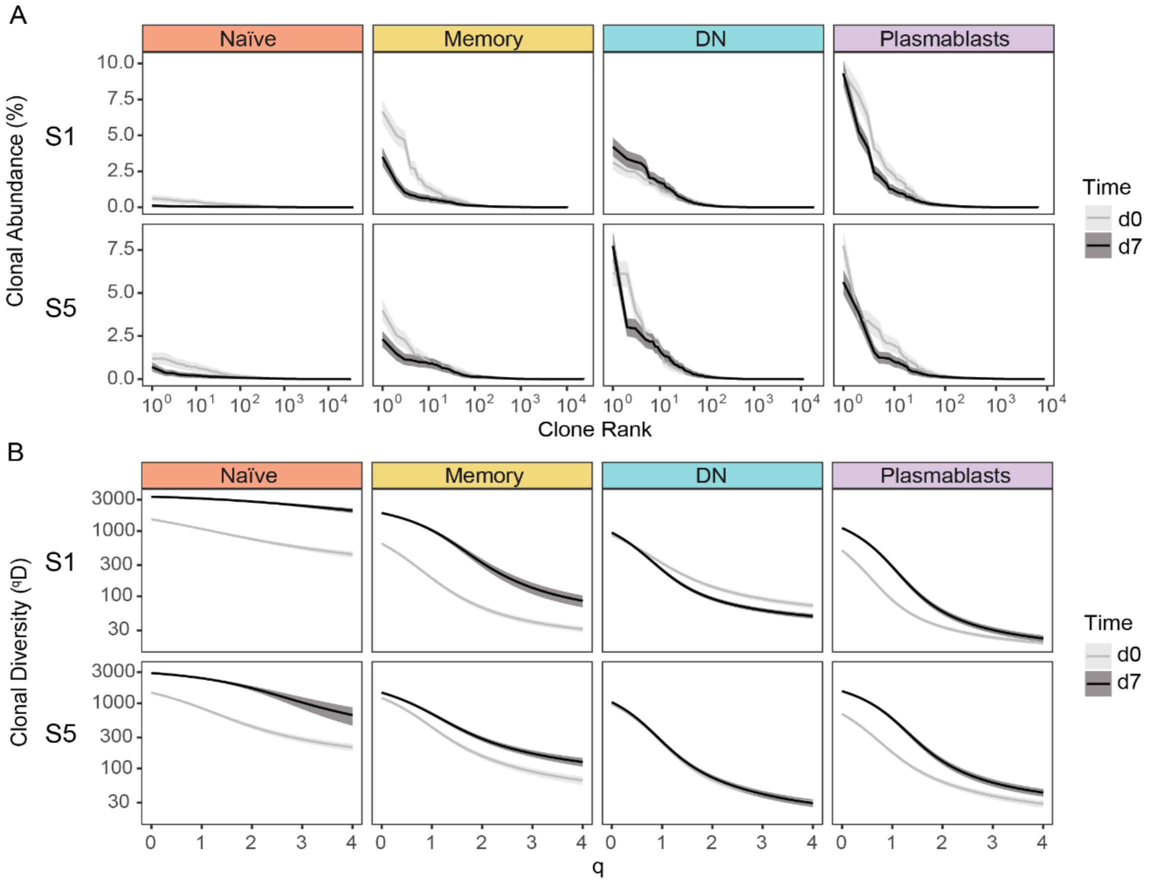


**Fig. S3:** Characteristics of the clonal repertoire of subjects 1 (S1) and 5 (S5), at day 0 before and day-7 after TBE vaccination. (A) Clonal abundance of the naïve, memory, DN and plasmablast repertoires. Clones were ranked according to decreasing sizes from left to right, and their relative abundance as percentage of the V(D)J sequence repertoire is represented. (B) Clonal diversity of the repertoire represented with Hill numbers ($^{q}D$) at different orders (q). For both clonal abundance and diversity calculations, a sampling of the unique sequence repertoire was performed, selecting N = 3906 unique sequences, with n = 1000 bootstrap iterations. This ensures the same number of sequences in each of the time points, and allows for correction of library sequencing depth correction. The gray area on both curves indicates the variability among the bootstrap samples.

**Supplementary Figure S4**

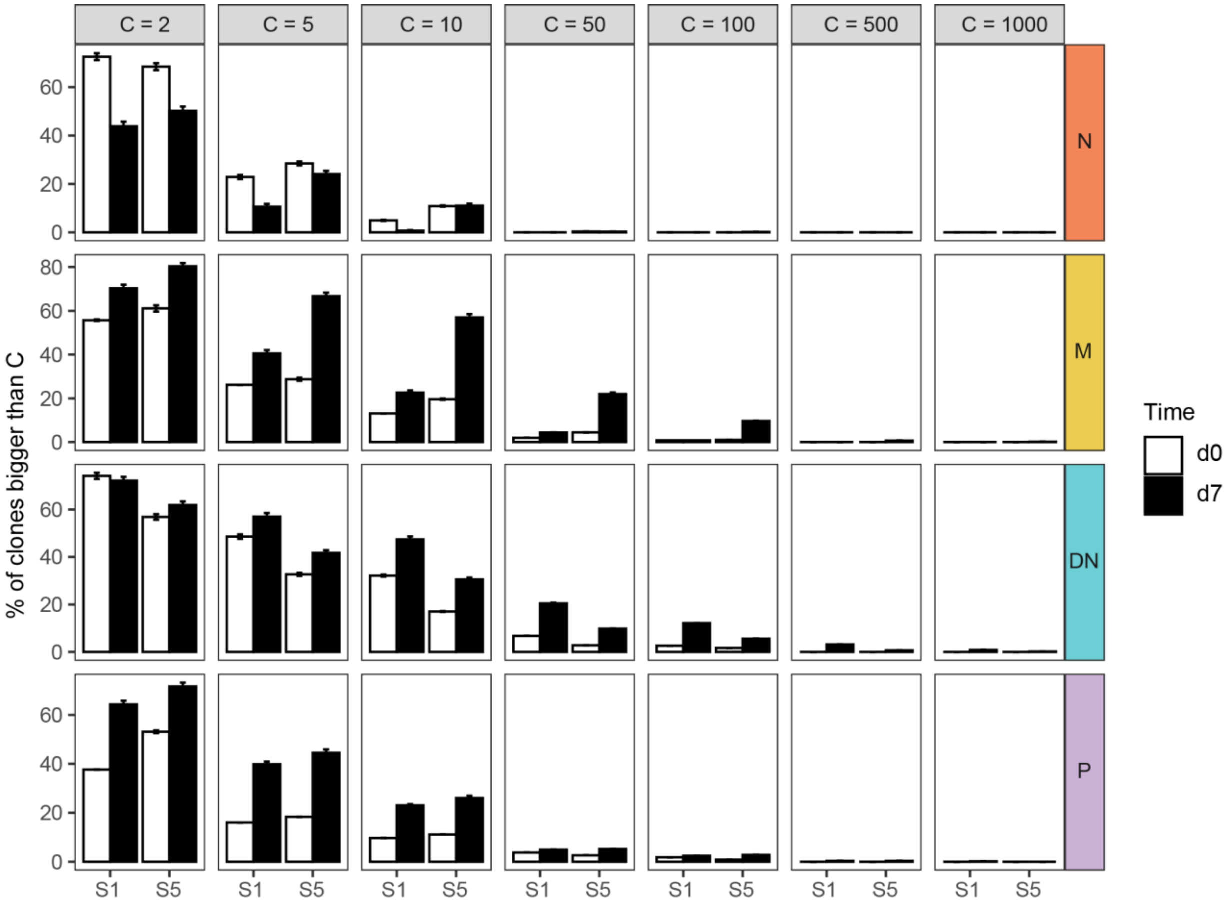


**Fig. S4:** Percentage of clones in the repertoire bigger or equal than size C, for different C values. To compute the percentages, a sampling of N=611 clones was performed according to their proportion in the repertoire. The error bars show the standard deviation of n=1000 bootstraps of the sampling.

**Supplementary Figure S5**

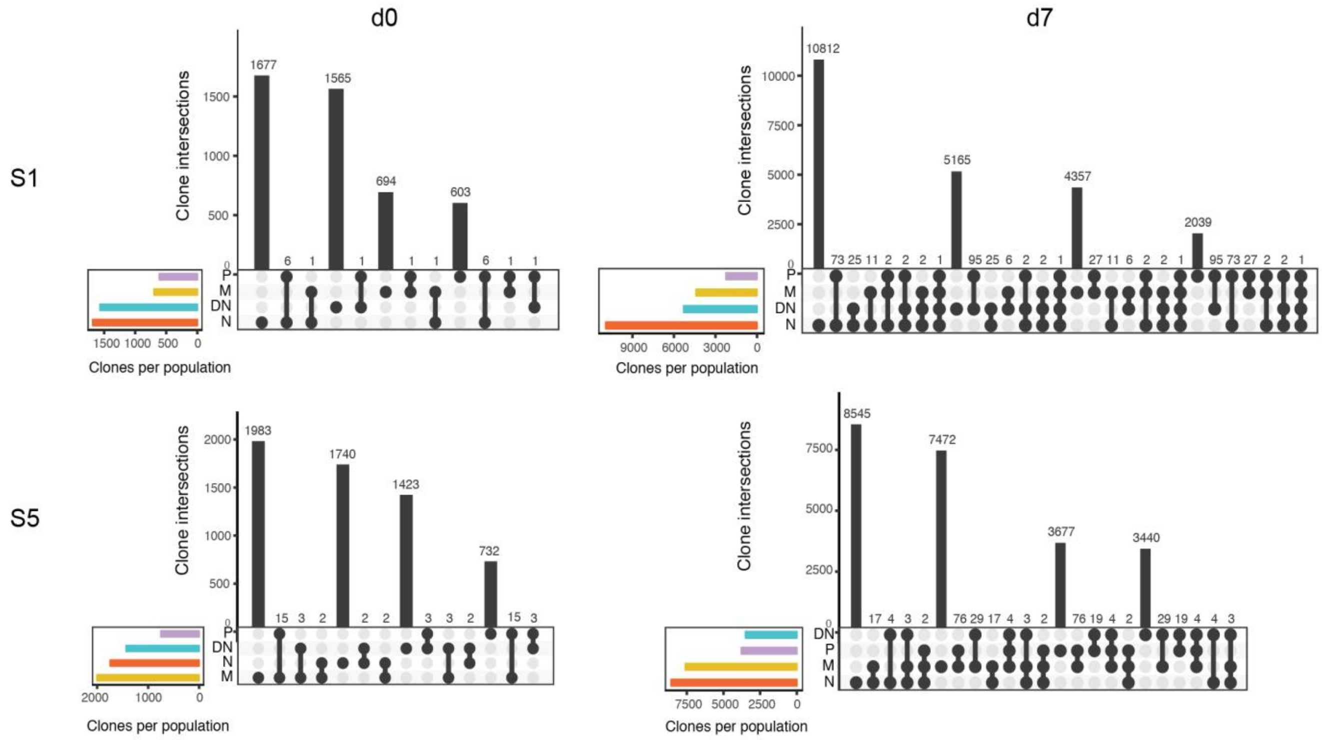

**Fig S5:** Upset plots representing the clonal overlaps among the B-cell populations of subjects 1 (S1) and 5 (S5) before and after vaccination (d7). Colored bars on the lower left for all the diagrams represent the total number of clones per population.

**Supplementary Table S1**. Genes of interest selected for the heatmap in Figure 2C. Adjusted p-values are indicated ($p < 0.05$, $p < 0.01$ or $p < 0.001$, ns = not significant) and the direction of regulation is indicated by colors (upregulated in green, down-regulated in red).

| SYMBOL | Probe set ID | DN vs N | DN vs PB | DN vs M | PB vs N | M vs N | PB vs M |
|---|---|---|---|---|---|---|---|
| **SLAMF7** | 16672654 | ns | P < 0.01 | ns | P < 0.001 | ns | P < 0.001 |
| **CR2** | 16676795 | ns | ns | ns | ns | ns | ns |
| **TRAF5** | 16677133 | ns | P < 0.001 | ns | P < 0.001 | ns | P < 0.001 |
| **MTOR** | 16681661 | ns | ns | ns | ns | ns | ns |
| **IFNLR1** | 16683551 | ns | ns | ns | ns | ns | ns |
| **FCRL5** | 16694886 | ns | ns | ns | ns | ns | ns |
| **CXCR5** | 16732037 | ns | P < 0.001 | ns | P < 0.001 | ns | P < 0.001 |
| **CD27** | 16747261 | 0 | P < 0.001 | 0 | P < 0.001 | P < 0.01 | 0 |
| **FOXO1** | 16778392 | ns | ns | ns | ns | ns | ns |
| **GPR183** | 16780592 | ns | ns | ns | ns | ns | P < 0.001 |
| **IGHD** | 16797395 | ns | ns | ns | P < 0.05 | ns | ns |
| **IGHD** | 16797425 | ns | ns | ns | ns | ns | ns |
| **IGHD** | 16797467 | ns | ns | ns | ns | ns | ns |
| **IL4R** | 16817254 | ns | P < 0.001 | ns | P < 0.001 | P < 0.01 | ns |
| **IL21R** | 16817271 | ns | ns | ns | ns | ns | ns |
| **CD19** | 16817489 | ns | ns | ns | P < 0.01 | ns | P < 0.001 |
| **ITGAX** | 16818272 | ns | ns | ns | ns | ns | ns |
| **IRF8** | 16821621 | ns | P < 0.01 | ns | P < 0.001 | ns | P < 0.001 |
| **TBX21** | 16835313 | ns | ns | ns | ns | ns | ns |
| **CCR7** | 16844381 | ns | P < 0.001 | ns | P < 0.001 | ns | P < 0.001 |
| **BCL2** | 16855673 | ns | ns | ns | ns | ns | ns |
| **ZEB2** | 16903356 | ns | ns | ns | ns | ns | ns |

| | | | | | | | |
|---|---|---|---|---|---|---|---|
| **IFIH1** | 16904365 | ns | ns | ns | ns | ns | ns |
| **IL10RB** | 16904365 | ns | ns | ns | ns | ns | ns |
| **BCL6** | 16962584 | ns | ns | ns | P < 0.001 | ns | ns |
| **CD38** | 16965268 | ns | P < 0.05 | ns | P < 0.01 | ns | P < 0.001 |
| **EBF1** | 17002278 | ns | P < 0.05 | ns | P < 0.001 | ns | P < 0.01 |
| **IRF4** | 17004167 | ns | ns | ns | P < 0.001 | ns | P < 0.001 |
| **PRDM1** | 17011279 | ns | P < 0.01 | ns | P < 0.001 | ns | P < 0.001 |
| **ABCB1** | 17059491 | ns | ns | ns | ns | ns | ns |
| **PAX5** | 17093973 | ns | P < 0.01 | ns | P < 0.001 | ns | P < 0.001 |
| **TLR7** | 17101531 | ns | ns | ns | ns | ns | ns |
| **CD24** | 17117110 | ns | P < 0.001 | ns | P < 0.001 | ns | P < 0.001 |
| **ZEB2** | 17117888 | ns | ns | ns | ns | ns | ns |
| **FCRL5** | 17120182 | ns | ns | ns | ns | ns | ns |

**Supplementary Table S2**: Basic repertoire characteristics of targeted immunoglobulin repertoire analysis at baseline and day-7 after vaccination including isotype frequency, VH family usage and mutational analysis

| | | **Vaccination subject 1** | | | | | | | |
|---|---|---|---|---|---|---|---|---|---|
| | | **Baseline** | | | | **Day 7** | | | |
| | | **N** | **M** | **DN** | **P** | **N** | **M** | **DN** | **P** |
| **Isotype Frequency** | *IgA %* | 0 | 26 | 15 | 26 | 0 | 26 | 44 | 35 |
| | *IgD %* | 77 | 0 | 0 | 0 | 2 | 0 | 0 | 0 |
| | *IgG %* | 0 | 41 | 84 | 48 | 0 | 48 | 52 | 48 |
| | *IgM %* | 23 | 33 | 1 | 27 | 98 | 25 | 4 | 17 |
| **VH-Family Usage** | *IGHV1 %* | 18 | 19 | 10 | 7 | 29 | 17 | 15 | 5 |
| | *IGHV2 %* | 5 | 2 | 5 | 17 | 25 | 17 | 5 | 31 |
| | *IGHV3 %* | 23 | 28 | 31 | 22 | 16 | 28 | 49 | 19 |
| | *IGHV4 %* | 44 | 38 | 26 | 22 | 15 | 27 | 25 | 23 |
| | *IGHV5 %* | 8 | 13 | 26 | 32 | 14 | 11 | 6 | 23 |
| **Mutation Analysis** | *Mean Mutation Count* | 6 | 21 | 16 | 24 | 7 | 21 | 24 | 21 |
| | *SD Mutation Count* | 10 | 11 | 12 | 9 | 9 | 12 | 14 | 11 |

| | | **Vaccination subject 5** | | | | | | | |
|---|---|---|---|---|---|---|---|---|---|
| | | **Baseline** | | | | **Day 7** | | | |
| | | **N** | **M** | **DN** | **P** | **N** | **M** | **DN** | **P** |
| **Isotype Frequency** | *IgA %* | 0 | 27 | 46 | 24 | 0 | 18 | 30 | 27 |
| | *IgD %* | 42 | 0 | 0 | 0 | 12 | 0 | 0 | 0 |
| | *IgG %* | 0 | 67 | 44 | 48 | 0 | 81 | 62 | 57 |
| | *IgM %* | 58 | 6 | 10 | 28 | 88 | 1 | 8 | 16 |
| **VH-Family Usage** | *IGHV1 %* | 28 | 11 | 27 | 16 | 21 | 27 | 28 | 15 |
| | *IGHV2 %* | 9 | 37 | 8 | 20 | 28 | 25 | 10 | 22 |
| | *IGHV3 %* | 30 | 7 | 36 | 13 | 20 | 6 | 24 | 17 |
| | *IGHV4 %* | 20 | 14 | 16 | 18 | 15 | 23 | 29 | 23 |
| | *IGHV5 %* | 13 | 30 | 13 | 33 | 15 | 18 | 9 | 23 |
| **Mutation Analysis** | *Mean Mutation Count* | 9 | 16 | 14 | 19 | 9 | 21 | 17 | 20 |
| | *SD Mutation Count* | 13 | 11 | 12 | 11 | 14 | 12 | 12 | 11 |

**Supplementary Table S3**: Tick borne encephalitis ELISA test of recombinant antibodies (rAb) and serum

| | **ELISA mean values** | **Interpretation** |
|---|---|---|
| **S1 day-0 (serum)** | 38 VIEU/ml | Negative |
| **S1 day-14 (serum)** | 352 VIEU/ml | Positive |
| **S5 day-0 (serum)** | 111 VIEU/ml | Borderline |
| **S5 day-14 (serum)** | 471 VIEU/ml | Positive |
| **S1 F4 day-9 (rAb)** | 138 VIEU/ml | Positive |
| **S1 C3 day-9 (rAb)** | 90 VIEU/ml | Borderline |
| **S5 D10 day-9 (rAb)** | Below detection limit | Negative |

*Negative < 63 VIEU/ml>; Borderline 63-126 VIEU/ml; Positive > 126 VIEU/ml*

**Supplementary Table S4**: Cell counts, number of VH sequences and number of clones

| | | **Day-0** | | | | **Day-7** | | | |
|---|---|---|---|---|---|---|---|---|---|
| | | *Cell counts* | *Unique sequences with 2 representatives* | *Number of sequences after Igblast* | *Number of clones* | *Cell counts* | *Unique sequences with 2 representatives* | *Number of sequences after Ig-blast* | *Number of clones* |
| S1 | N | 65000 | 5818 | 4128 | 1684 | 350000 | 53409 | 21614 | 10928 |
| | M | 15000 | 5148 | 4441 | 696 | 310000 | 18931 | 16233 | 4406 |
| | DN | 9163 | 16317 | 11512 | 1566 | 48000 | 70578 | 53190 | 5296 |
| | P | 1009 | 6124 | 5221 | 611 | 1396 | 14417 | 12678 | 2239 |
| S5 | N | 82000 | 8656 | 4887 | 1744 | 170000 | 34893 | 18594 | 8571 |
| | M | 61000 | 9528 | 8098 | 2003 | 110000 | 65015 | 55300 | 7603 |
| | DN | 25000 | 10800 | 7370 | 1431 | 84000 | 28797 | 23288 | 3499 |
| | P | 6251 | 4712 | 4209 | 750 | 10000 | 19654 | 17356 | 3778 |

S1 = subject 1; S5 = subject 5
